# Beyond Pairwise Connections in Complex Systems: Insights into the Human Multiscale Psychotic Brain

**DOI:** 10.1101/2025.03.18.643985

**Authors:** Qiang Li, Shujian Yu, Jesus Malo, Godfrey D. Pearlson, Yu-Ping Wang, Vince D Calhoun

## Abstract

Complex biological systems, like the brain, exhibit intricate multiway and multiscale interactions that drive emergent behaviors. In psychiatry, neural processes extend beyond pairwise connectivity, involving higher-order interactions critical for understanding mental disorders. Conventional brain network studies focus on pairwise links, offering insights into basic connectivity but failing to capture the complexity of neural dysfunction in psychiatric conditions. This study aims to bridge this gap by applying a matrix-based entropy functional to estimate total correlation, a mathematical framework that incorporates multivariate information measures extending beyond pairwise interactions. We apply this framework to fMRI-ICA-derived multiscale brain networks, enabling the investigation of interactions beyond pairwise relationships in the human multiscale brain. Additionally, this approach holds promise for psychiatric studies, providing a new lens through which to explore beyond pairwise brain network interactions. By examining both triple interactions and the latent factors underlying the triadic relationships among intrinsic brain connectivity networks through tensor decomposition, our study presents a novel approach to understanding higher-order brain dynamics. This framework not only enhances our understanding of complex brain functions but also offers new opportunities for investigating pathophysiology, potentially informing more targeted diagnostic and therapeutic strategies. Moreover, the methodology of analyzing multiway interactions beyond pairwise connections can be applied to any signal analysis. In this study, we specifically explore its application to neural signals, demonstrating its power in uncovering complex multiway interaction patterns of brain activity, which provide a window to explore connectivity beyond pairwise interactions in the multiscale functionality of the brain.

## I Introduction

Accessing interactions beyond pairwise in complex systems is essential for understanding their collective behavior, as these interactions provide crucial insights that go beyond simple pairwise relationships [1]. The human brain is organized in a hierarchical, multiway, and multiscale structure, which facilitates efficient information processing and is crucial for maintaining functional interactions across diverse brain networks [2, 3]. Functional connectivity provides a valuable window through which we can measure the interactions between these brain networks [4]. While traditional approaches have primarily focused on pairwise interactions, understanding beyond pairwise relationships is crucial for capturing the full complexity of the brain’s dynamic networks [5–9]. These beyond pairwise interactions offer deeper insights into the brain’s intricate connectivity, revealing how multiple brain networks interact synergistically or redundantly. By exploring these higher-order interactions, we can gain a more refined understanding of the brain’s organizational structure and its underlying mechanisms, both in normal and pathological states [1, 10–12].

To estimate beyond pairwise connectivity, numerous metrics have been applied in brain network studies, uncovering interesting findings that are often missed by pairwise metrics. From an information-theoretical standpoint, total correlation [13] and dual total correlation [14] are two metrics that can be used to assess interactions beyond pairwise relationships. These metrics have been employed to investigate higher-order interactions in both healthy brains [5, 7] and in various brain disorders [8, 10]. They uncover connections that are frequently overlooked in pairwise analyses. From a graph theory standpoint, hypergraphs provide a crucial framework for understanding higher-order interactions within graphs. When combined with graph neural networks, they have been increasingly applied in brain network science research [15]. From the perspective of topology and geometric data analysis, methods such as persistent homology, Euler characteristic, tensor decomposition, and curvature have been utilized to quantify higher-order interactions [16, 17].

In studies of functional connections via resting-state functional MRI (rsfMRI), common metrics such as Pearson correlation and mutual information are used to evaluate both linear and nonlinear aspects of pairwise functional connectivity [5, 6, 18]. However, these methods are limited because they only account for simple pair-wise interactions, ignoring the complex, multiway interactions that characterize brain dynamics [5, 8, 10–12]. To overcome these limitations, an information theory-based measure called total correlation [13, 19–21] can be used to detect more complex, beyond-pairwise interactions. Studies have shown that total correlation not only provides richer insights into brain connectivity but also improves diagnostic accuracy for brain disorders [1, 7, 10, 22]. Furthermore, this study adopts a more adaptive strategy, utilizing data-driven methods like ICA to pinpoint functionally coherent regions unique to each subject [23, 24]. The ICA identifies intrinsic connectivity networks (ICNs) directly from the BOLD signal, removing the reliance on pre-defined anatomical or atlas-based divisions [25–27]. This approach allows for the discovery of brain networks based solely on the observed data.

To tackle the challenges of beyond-pairwise dependencies, we propose an approach that utilizes matrix-based rényi’s entropy functional to generate beyond-pairwise descriptors. Specifically, this method focuses on estimating higher-order information interactions to gain deeper insights into the complex interdependencies within the human brain. By analyzing how brain connectivity evolves as networks are introduced, we can gain a deeper understanding of how different networks interact, including their redundant relationships. This approach enables us to clarify the role of each network, providing a more comprehensive understanding of brain function and advancing psychiatric brain studies to uncover potential biomarkers. Furthermore, our findings underscore the significant potential of beyond-pairwise interactions for applications in other areas of network science and the analysis of different signal modalities.

## II Results

### A Multiway Interactions at Multiple Scales

Capturing multi-way interactions in the human brain across multiple scales is essential for understanding information exchange within the brain. The number of interactions grows as 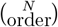, where *N* represents the total number of neurons, synapses, or networks, and the order refers to the interaction size (e.g., 2-way, 3-way, 4-way, etc.). With 100 billion neurons (*N* = 1.0000 × 10^11^) or 100 trillion synapses (*N* = 10^14^), the number of interactions increases astronomically as the order of interactions rises (i.e., for order = 2, 3, 4, …), as shown in ***Fig. 1A***.

**Fig 1:**
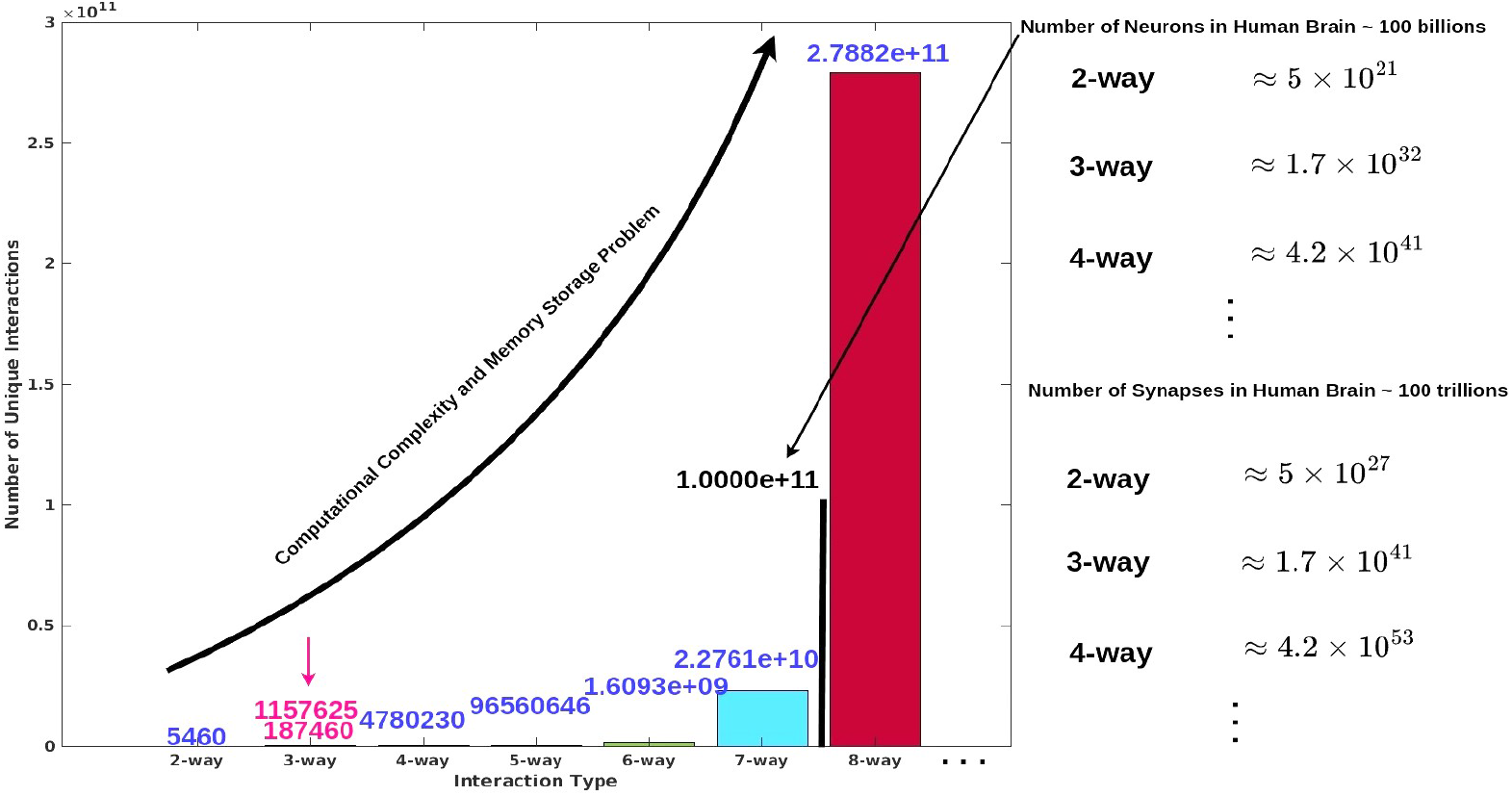
Computational Complexity of Multiway Interactions in the Human Brain. At the microscopic level, the human brain contains approximately 100 billion neurons. When considering multiway interactions (i.e., 2-way, 3-way, 4-way interactions, and so on), the number of interactions increases exponentially. If we extend this to synaptic interactions, the number of interactions increases sharply, and current computational infrastructure cannot meet the demands for computing and storage (at least as of 2025). At the macroscopic level, we consider 105 brain networks. Even with multiway interactions at this scale, we still face significant computational and storage challenges. To balance these needs, we ultimately decided to focus on 3-way (triple) interactions to explore higher-order interactions in both normal and psychotic brains.

As seen, the computational complexity for calculating 2-way, 3-way, and 4-way interactions grows exponentially with *N* at the microscopic level. As the number of interactions increases, the memory required to store these combinations becomes far beyond the capacity of typical hardware, especially for very large *N*. Managing such large-scale computations poses significant computational and storage challenges, and current computers are unable to handle this burden effectively.

In our study, we focus on the macroscopic level of brain network interactions, considering 105 networks that collectively cover the entire brain [28, 29]. The number of interactions grows rapidly with the order of the interaction: for 2-way interactions 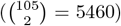, 3-way interactions 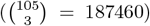, 4-way interactions 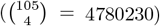, 5-way interactions 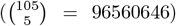 6-way interactions 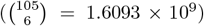, 7-way interactions 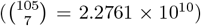, and 8-way interactions 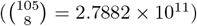, and so on. Given the computational complexity, the challenges of accurately estimating interactions, and the difficulty of visualizing higher-order interactions, we have decided to focus on 3-way (triple) interactions for our analysis.

### B Beyond Homogeneous: Multiscale Brain

The homogeneous brain treats each network as uniform, whereas a multiscale brain captures its complexity by analyzing networks at different spatial resolutions, which is essential for understanding the diverse and dynamic nature of brain connectivity. To extract ICNs from our resting-state data, we utilized the new multiscale *NeuroMark fMRI 2.2* template (available for download at https://trendscenter.org/data/), a multiscale brain network template derived from over 100K subjects [28, 29]. This template contains 105 networks extracted across various spatial resolutions, as shown in ***Fig. 2A***. These ICNs were derived from over 20 different studies and processed using a group multi-scale ICA approach with 8 distinct model orders [28]. Higher model orders typically correspond to higher spatial resolution, while lower model orders integrate features across larger brain regions. By incorporating data from multiple scales, we can model a more diverse range of intrinsic connectivity networks.

**Fig 2:**
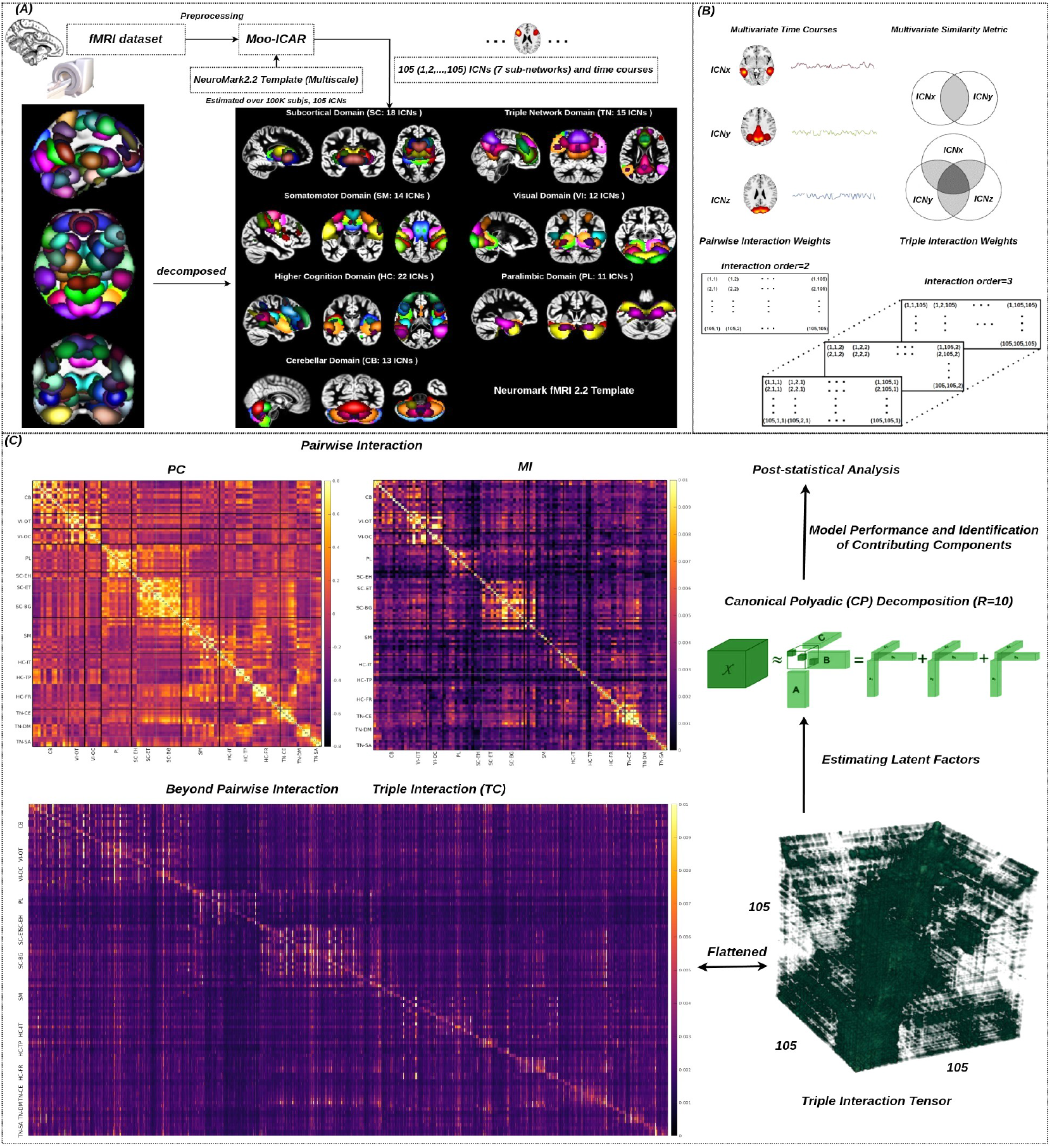
Flowchart for Analyzing Beyond-Pairwise Multiscale Interactions Within Brain Networks. The preprocessed resting-state fMRI data were input into the Multivariate Objective Optimization ICA with Reference (MOO-ICAR) using the spatially constrained *NeuroMark2.2 Template*. This process generated subject-specific estimates of 105 intrinsic connectivity networks (ICNs) and their corresponding time courses in resting-state fMRI. These 105 ICNs were subsequently grouped into seven large brain domains: visual (VI), cerebellar (CB), subcortical (SC), somatomotor (SM), paralimbic (PL), higher cognitive (HC), and the triple network domain (TN), as shown in ***A***. As brain networks are inherently interactive, with information exchange occurring both pairwise and beyond pairwise, we estimated both pairwise (interaction order=2) and triple interactions (interaction order=3) among the ICNs, providing insights into multiway interactions within the brain, as illustrated in ***B, C***. To uncover hidden latent factors in triple interaction tensors, we applied Canonical Polyadic Decomposition with Alternating Least Squares (CP-ALS) [30] to the tensor, decomposing it into 10 components. We then evaluated the model’s fitting performance and conducted post-statistical analysis on the latent factors, as shown on the right side of ***C***.

The 105 ICNs are organized into 14 major functional domains, including: visual domain (VI, 12 sub-networks; occipitotemporal subdomain (OT) and occipital subdomain (OC)), cerebellar domain (CB, 13 sub-networks), sub-cortical domain (SC, 18 sub-networks; extended hippocampal subdomain (EH), extended thalamic subdomain (ET), and basal ganglia subdomain (BG)), sensorimotor domain (SM, 14 sub-networks), high cognition domain (HC, 22 sub-networks; insular-temporal subdomain (IT), temporoparietal subdomain (TP), and frontal subdomain (FR)), triple network domain (15 sub-networks; central executive subdomain (CE), default mode subdomain (DM), and salience subdomain (SA)), and paralimbic domain (PL, 11 sub-networks).

### C Beyond Pairwise: Triple Interactions

If we only consider pairwise interactions, there are 105^2^ = 5460 pairwise interactions within the multiscale brain network, calculated based on the widely used metrics of Pearson correlation and mutual information, as shown in ***Fig. 2B,C***. These pairwise interactions offer an initial understanding of the pairwise relationships between different brain regions, capturing both linear and nonlinear functional dependencies. When considering higher-order interactions, specifically for ***k*** = 3, the number of possible triple interactions increases dramatically, amounting to 105^3^ = 1, 157, 625 interactions, and there are 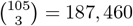 unique sets of triple interactions, as illustrated in ***Fig. 2B,C***. It provides a more comprehensive framework for examining the intricate and multidimensional nature of connectivity within multiscale human brain networks. These interactions allow for a deeper exploration of the complex, higher-order relationships that underlie brain function, which are crucial for understanding the brain’s dynamic and evolving connectivity patterns across different scales.

Compared to pairwise interactions, the number of triple interactions increases by a factor of 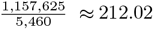 if all triple interactions are considered, and the number of triple interactions increases by a factor of 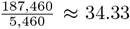 if all unique triple interactions are considered. The computation of all triple interactions for each subject is highly computationally intensive and requires significant time, even when utilizing large cluster servers. In our case, we used parallel computing using 8 GTX 1080Ti GPUs, and the entire process took approximately one month to complete.

In summary, considering higher-order relationships within the multiscale brain network leads to a substantial increase in complexity. This growth underscores the importance of incorporating triple interactions, as they reveal more intricate dependencies and provide a richer understanding of the brain’s functional architecture.

### D Triple Interactions Preserve Meaningful Brain Connectivity Patterns

The triple interactions are represented as a 3D tensor, as shown in ***Fig. 3A***, which illustrates the interactions between triple ICNs. Different thresholds were applied to the original tensor, revealing clear clustering patterns along the diagonal. To visualize these triple functional connectivity patterns more directly, we flattened the 3D tensor into a 2D matrix, where the connectivity pattern is again observed along the diagonal, as shown in ***Fig. 3B***. Compared to pairwise functional connectivity, derived from Pearson correlation and mutual information (shown on the left side of ***Fig. 2C***), we demonstrate that meaningful brain network connectivity is captured within triple network interactions. Given the large number of triple interactions, we identified the strongest and weakest interactions within the flattened triple interaction matrix. The strongest unique triple interaction involves HC-IT (ICN 70), PL (ICN 34), and VI-OT (ICN 15), as depicted on the left side of ***Fig. 3C***. In contrast, the weakest triple interaction involves SC-EH (ICN 38), PL (ICN 36), and SM (ICN 59). This is a straightforward way to understand the connections within these large, complex tensors, while another approach involves extracting the latent factors that underlie them.

**Fig 3:**
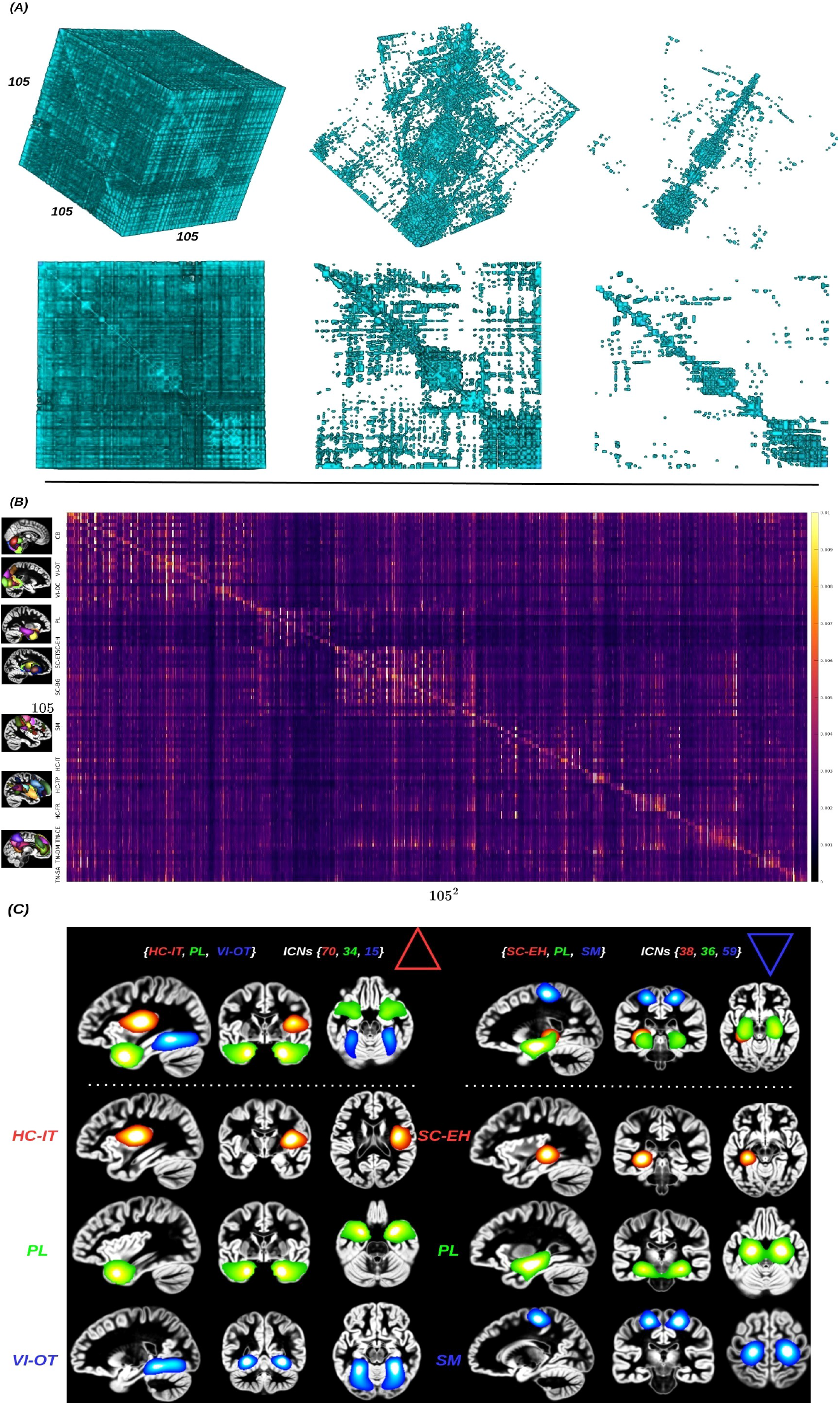
Multiscale Triple Interactions Across Brain Networks. To enhance the visualization of cluster patterns in the diagonal of the multiscale triple interaction tensor, different thresholds were applied. The results are displayed from left to right in the first row, as shown in ***A***. Furthermore, varying viewing angles in the second row emphasize the internal structure of the tensor. The 3D tensor was then flattened into a 2D matrix, revealing the internal structure more distinctly, as shown in ***B***. There are a total of 105^3^ triple interactions, with the maximum (HC-IT, PL, VI-OT) and minimum (SC-EH, PL, SM)interactions identified within the multiscale human brain, as shown on the left and right sides of ***C***.

### E Identified Latent Components in Triad Connectivity Tensor Through Canonical Polyadic (CP) Decomposition

To uncover hidden latent factors in complex triadic interactions, we applied the CP decomposition to triadic interaction tensors with a rank of 10, as shown in ***Fig. 4A***. The performance of the CP model fitting was evaluated and is presented in ***Fig. 4B***. The dot plots below show the reconstruction errors for 30 model fits (in blue) and the reconstruction errors for the true latent factors (in red). Our results demonstrate that the CP model fitting produced minimal errors and achieved stable, high-quality performance. Meanwhile, an illustration of a 10-component nonnegative factorization is shown in ***Fig. 4C***. We plot each component as a row in the figure. The component numbers are displayed along the left side of the plot. This plot is a scatter chart that visualizes the activity of the ICNs.

**Fig 4:**
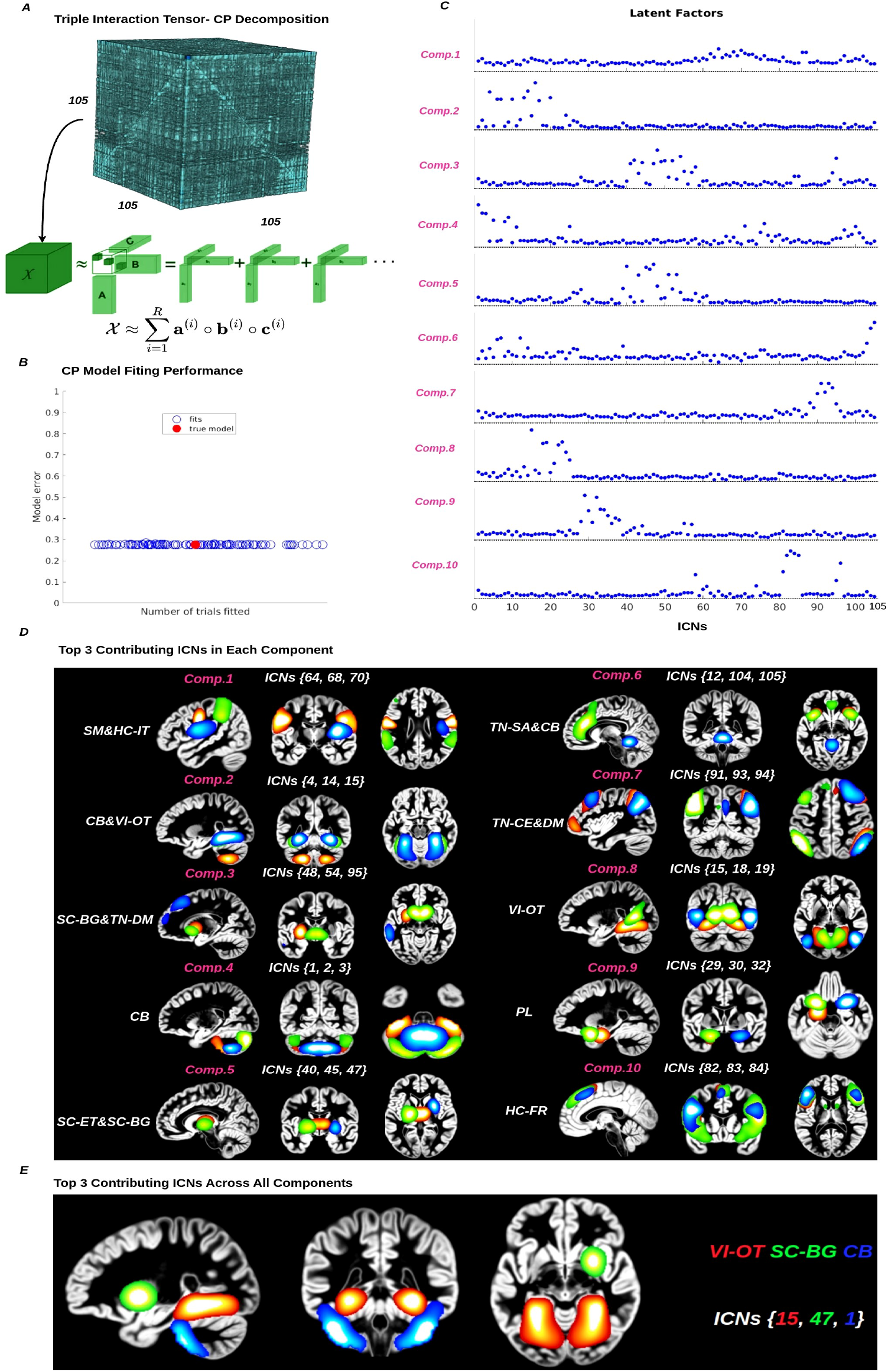
Latent Factors in Triadic Interactions via CP Decomposition. The CP decomposition was applied to the triple interaction tensor to explore latent factors, revealing hidden patterns within the interactions, as shown in ***A***. In our case, we used a rank of *R* = 10 to decompose the triple interaction into 10 components. The model’s fitting performance was assessed, as shown in ***B***, and the resulting latent factors are displayed in ***C***. From these decomposed components, we selected the top three contributing ICNs for each component, as illustrated in ***D***. The top 3 most contributing ICNs across all components were identified, with the three strongest interactions in the human brain highlighted, as shown in ***E***.

Furthermore, several observations can be made from the decomposed latent factors, as shown in ***Fig. 4D***. Firstly, component 1 remains consistently active across all ICNs. The top three ICNs, SM (ICNs 64 and 68) and HC-IT (ICN 70), stand out compared to the activity of other ICNs. Component 2 separates the ICN targets, with the largest magnitude associated with this component. The top three ICNs with the highest values are CB (ICN 4) and VI-OT (ICNs 14&15). Component 3 separates the ICNs, with the top three ICNs having the highest values: SC-BG (ICNs 48&54) and TN-DM (ICN 95). Component 4 separates the ICNs, with the top three ICNs having the largest magnitudes: CB (ICNs 1&2&3). Component 5 separates the ICNs, with the top three ICNs being SC-ET (ICNs 40&45) and SC-BG (ICN 47). Component 6 separates the ICNs, with the top three ICNs being TN-SA (ICNs 104&105) and CB (ICN 12). Component 7 separates the ICNs, with the top three ICNs being TN-CE (ICNs 91&93) and DM (ICN 94). Component 8 separates the ICNs, with the top three ICNs being VI-OT (ICNs 15&18&19). Component 9 separates the ICNs, with the top three ICNs being PL (ICNs 29&30&32). Component 10 separates the ICNs, with the top three ICNs being HC-FR (ICNs 82&83&84). Finally, we also examined the top three contributing ICNs in triple interactions across all components: VI-OT (ICN 15), SC-BG (ICN 47), and CB (ICN 1), as illustrated in ***Fig. 4E***.

In summary, we observed that the TN networks play a major role in complex triadic interactions, which aligns with existing neural mechanisms [31]. The TN network domains, including DM, SA, and CE, contribute significantly, with DM in particular playing a dominant functional role during resting state. Additionally, we found that other networks, such as VI-OT, HC-IT/FR, CB, SC-BG/ET, SM, and PL, also make important contributions to these interactions. This suggests that large-scale networks engage in higher-order interactions to regulate brain functions.

These findings are further supported by the previous analysis [32], which identified 10 stable and consistent patterns with potential functional relevance in resting states. These patterns encompass regions involved in motor function, visual processing, executive functioning, auditory processing, memory, and the default-mode network, each exhibiting BOLD signal changes of up to 3% [32]. These results also align with our findings, further strengthening the validity of our conclusions. Overall, our results are compelling, as they not only effectively separate the ICNs but also uncover additional activities that offer insightful and meaningful interpretations. Together, these findings suggest that large-scale networks play a crucial role in complex, higher-order brain interactions and contribute to the dynamic regulation of brain functions.

### F Identifying Aberrant Networks in Schizophrenia Through Triple Interaction Analysis

To explore triple interactions in the psychotic brain, we examined triple connections in schizophrenia (SZ). First, we identified the most significant ICNs using a threshold of 0.3 from each CP decomposition component in SZ compared to the control group, as illustrated in ***Fig. 5A***. The most significant deviations in ICNs were found in components 2, 4, 6, 7, 8, 9, and 10, while no significant ICNs were observed in components 1, 3, and 5 under the 0.3 threshold.

**Fig 5:**
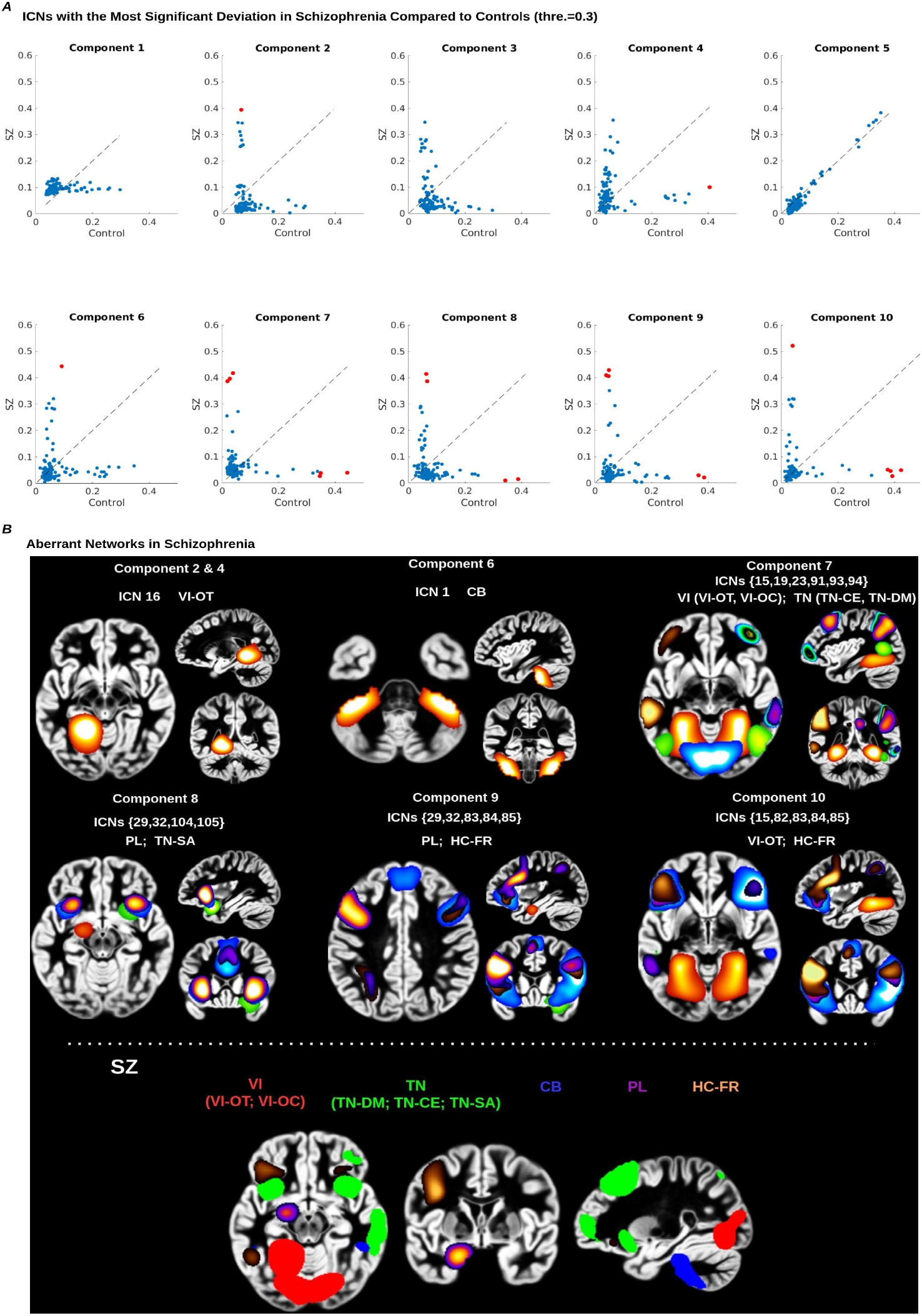
Aberrant Networks in Schizophrenia. The most significant deviations of ICNs in SZ compared to controls were identified using a magnitude threshold of 0.3 across all components from CP decompositions and labeled in red, as shown in ***A***. The spatial map of each identified deviated ICN was presented, and finally, the most aberrant networks in SZ were identified, including VI (VI-OT; VI-OC), TN (TN-DM, CE, SA), CB, PL, and HC-FR, as shown in ***B***.

To more precisely identify aberrant networks in SZ, we analyzed the most consistently affected significant networks across all deviated ICNs in each component. We found that VI (VI-OT; VI-OC), TN (TN-DM, TN-CE, TN-SA), CB, PL, and HC-FR showed the most significant deviations compared to normal brains, as shown in ***Fig. 5B***. In summary, this provides a clear identification of several networks affected in SZ, which could serve as potential biomarkers for SZ treatment. Furthermore, it opens new avenues for understanding brain disorders through the lens of high-order interactions.

## III Discussion

In this study, we explore beyond pairwise interactions in the human multiscale brain by estimating triple functional connectivity through total correlation [13, 19]. This approach not only captures higher-order interactions but also preserves meaningful functional connectivity patterns, offering a more nuanced understanding compared to traditional pairwise functional connectivity methods. To delve deeper into the hidden latent factors behind these complex large-scale triple interactions, we applied tensor decomposition to the triple interaction tensors. This allowed us to uncover latent factors that reveal the underlying structure of these interactions. By analyzing these factors, we gain valuable insights into which components contribute more or less to the overall complexity of the triple interactions.

Additionally, we extended this pipeline to investigate psychotic brain conditions, specifically schizophrenia, and identified several networks (i.e., visual, triple, cerebellar, paralimbic, and frontal networks) that exhibit significant changes in schizophrenia compared to normal controls. These identified networks may serve as potential biomarkers for schizophrenia treatment, offering new avenues for diagnosis and therapeutic strategies. In numerous previous studies, each of these networks has been identified as being involved in schizophrenia separately [22, 33–35]. However, by exploring latent factors in triple interactions, we are able to identify all of these aberrant networks together. In summary, the exploration of beyond pairwise interactions provides a fresh perspective on studying higher-order interactions within the complex brain, revealing new insights into both normal and altered brain dynamics. Moreover, our method holds great potential for application in other brain disorders, network science, and even the analysis of other modalities of signals.

When considering computational complexity, our focus on triple interactions in the human multiscale brain has provided valuable new insights. However, an equally important aspect to consider is the complexity of interactions beyond triple interactions [36]. The brain’s connectivity involves far more than just triplets, with a rich and intricate network of interactions occurring across multiple levels. Extending the analysis to include interactions beyond triples could reveal previously overlooked aspects of brain dynamics, offering novel perspectives and insights that may have remained unnoticed in previous studies. Future research that explores these higher-order interactions will likely provide deeper understanding and potentially uncover new dimensions of brain function and dysfunction. Additionally, it is crucial to recognize that higher-order interactions in the human brain are dynamic rather than static, a factor that has significant implications for both the interpretation of our results and the broader understanding of brain connectivity [36, 37]. By examining how interactions between brain networks evolve over time, dynamic high-order functional connectivity approaches can reveal transient states and fluctuations that static models are unable to capture [38].

To explore the hidden latent factors, we applied CP decomposition [30, 39, 40] to the triple interaction tensors, which provided valuable insights into the underlying structure of these large and complex interactions. However, CP decomposition does have limitations when handling large tensors, and alternative tensor decomposition methods may be worth exploring [30, 39, 40]. By using other decomposition techniques, we could obtain more precise latent factors, which would ultimately enhance our ability to identify more accurate biomarkers for brain disorders.

In summary, while analyzing triple interactions provides valuable insights into both human and psychotic brain connectivity offering a deeper understanding beyond what pairwise interactions can reveal, it also introduces significant challenges. These challenges reflect the broader complexities inherent in studying multiway interactions, highlighting the need for advanced methods to fully capture the intricacies of brain dynamics.

## IV Methodology

### A rsfMRI Dataset

Considering the computational complexity and memory challenges, we analyzed resting-state fMRI data from 164 subjects, including 114 unrelated normal controls (NC), all from the multi-site Bipolar and Schizophrenia Network on Intermediate Phenotypes study [41, 42]. The scanning period was approximately five minutes across all sites. All subjects were psychiatrically stable and on stable medication regimens at the time of the study. Participants were instructed to rest with their eyes closed while remaining awake. Detailed scanning information for the entire study sample is provided elsewhere [41]. To further explore the validity of beyond pairwise interactions, we extended our methods to study brain disorders and used data from 50 typical schizophrenia subjects in this analysis.

### B rsfMRI Dataset Processing

The rigorous preprocessing pipeline applied to our resting-state fMRI (rsfMRI) data encompasses several essential steps designed to ensure the integrity and reliability of the data for subsequent analysis. First, careful quality control procedures were applied to identify and retain high-quality data, thereby ensuring the reliability of our analyses. Next, each participant’s rsfMRI data underwent a standardized preprocessing pipeline, which included rigid body motion correction, slice timing correction, and distortion correction. The preprocessed data were then registered to a common spatial template, resampled to isotropic voxels of 3mm^3^, and spatially smoothed with a Gaussian kernel having a full-width at half-maximum of 6mm.

### C Spatially Constrained ICA on rsfMRI

A spatially constrained ICA (scICA) approach, specifically the Multivariate Objective Optimization ICA with Reference (MOO-ICAR), was implemented using the GIFT software toolbox (http://trendscenter.org/software/gift) [43]. The MOO-ICAR method estimates subject-level independent components (ICs) by leveraging existing network templates as spatial references [25, 33, 37, 43, 44]. One of its key advantages is maintaining consistent correspondence between estimated ICs across subjects. Furthermore, the scICA framework allows for the customization of the network template used as a spatial reference during the ICA decomposition. This flexibility supports both disease-specific network analyses and more generalized evaluations of well-established functional networks, making it suitable for diverse populations [23, 25, 28, 45–47].

The MOO-ICAR algorithm, which implements scICA, optimizes two objective functions: one that enhances the overall independence of the networks, and another that improves the alignment of each subject-specific network with its corresponding template [25]. The two objective functions, 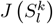 and 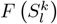, are outlined in the following equation, which illustrates how the *l*^*th*^ network can be estimated for the *k*^*th*^ subject using the network template *S*_*l*_ as a reference:

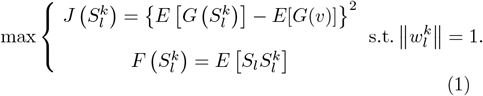

In this formulation, 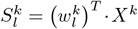 represents the estimated *l*^*th*^ network for the *k*^*th*^ subject, where *X*^*k*^ is the whitened fMRI data matrix of the *k*^*th*^ subject, and 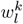 is the unmixing column vector, which is solved in the optimization functions. The objective function 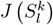 optimizes the independence of 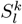 using negentropy. Here, *v* is a Gaussian variable with zero mean and unit variance, *G*(.) is a nonquadratic function, and *E*[.] denotes the expectation of the variable. The function 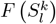 optimizes the correspondence between the template network 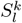 and the subject-specific network *S*^*k*^. The optimization problem is solved by combining the two objective functions through a linear weighted sum, with each weight set to 0.5. By applying scICA with MOO-ICAR to each scan, subject-specific ICNs are obtained for each of the N network templates, along with their associated time courses.

In this study, we used the *NeuroMark fMRI 2.2 template* (available for download at https://trendscenter.org/data/) along with the MOO-ICAR framework for scICA on rsfMRI data. It enabled us to extract subject-specific ICNs and their associated time courses. This template includes 105 high-fidelity ICNs identified and reliably replicated across datasets with over 100K subjects [28, 29].

### D Multivariate Information Measures of the Human Brain Using Matrix-Based Entropy Functional

#### 1 Shannon Information

The *Shannon entropy* **H**(*X*), or simply entropy, of a continuous random variable (RV) *X* ∈ 𝒳 with probability density function *f*_*X*_, its differential entropy is defined as,

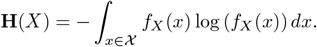

Then, the *Mutual Information* between *X* and another continuous RV *Y* ∈ 𝒴 is given by,

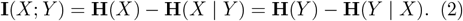

The *Mutual Information* measures the dependency between *X* and *Y*, and attains its minimum, equal to zero, if they are independent.

#### 2 Renyi’s α Entropy Functional

In information theory, a natural extension of the well-known *Shannon’s entropy* [48] is the *Renyi’s α entropy* [49]. For a random variable *X* with probability density function *p*(*x*) in a finite set 𝒳, the *α* entropy is defined as:

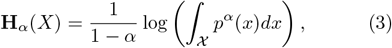

with *α ≠* 1 and *α* ≥0. In the limiting case where *α* →1, it reduces to *Shannon’s entropy* [50].

#### 3 Matrix–Based Information Estimator

In practice, given *m* realizations sampled from *p*(*x*), i.e., 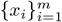, Sanchez Giraldo *et al*. [51] suggests that one can evaluate **H**_*α*_(*X*) without estimating *p*(*x*). Specifically, the so-called *matrix-based Renyi’s α entropy* is given as follows:

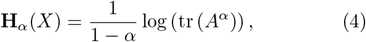

where *A* ∈ ℝ^*m*×*m*^ is a (normalized) Gram matrix with elements *A*_*ij*_ = *K*_*ij*_*/*tr(*K*), *K*_*ij*_ = *κ* (*x*_*i*_, *x*_*j*_) in which *κ* stands for a positive definite and infinitely divisible kernel such as Gaussian. tr(*) refers to matrix trace. As in [19], we set *α* = 1.01 to approximate Shannon entropy and choose a Gaussian kernel *G*_*σ*_ with width *σ*, given by,

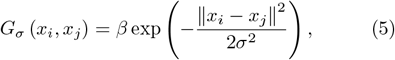

where *β* is a constant whose value is irrelevant because it is canceled out in the normalized Gram matrix.

#### 4 Bivariate Dependencies from *Mutual Information*

Bivariate dependencies can be inferred from mutual information, and if two variables are directly dependent, then the values are zeros. Given Eqs. 2 and 4, the *Renyi’s α entropy mutual information* **I**_*a*_(*X*^1^, *X*^2^) between variables *X*^1^ and *X*^2^ in analogy of *Shannon’s mutual information* is given by:

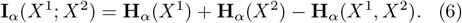

*Mutual information* is the most widely used metric for quantifying statistical dependency between two variables [50], but it omits higher-order interactions in the system; it does not capture all global information in the system.

#### 5 Kernel Width Selection

The value of the kernel width *σ* is central to the performance of the estimator described in Eq. 5. The following properties hold for the Gaussian kernel:

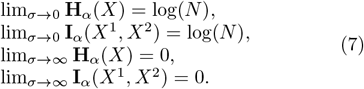

They imply that the value of *σ* controls the operating point of the estimator relative to the bounds because a value too large or too small saturates **H**_*α*_(*X*) and **I**_*α*_(*X*^1^, *X*^2^) to 0 and log(*N*), respectively. This saturation has to be avoided to have discriminative estimates. Therefore, a suitable value of *σ* has to be determined for an RV *X* of d dimensions and *N* samples.

A common rule for the Gaussian kernel is Silverman’s rule of thumb [52] that comes from the literature of density estimation. For the *j*-th dimension of *X*, it is given by

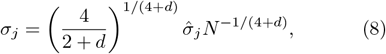

where 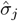 is the empirical standard deviation of the *j*-th dimension. In our study, the value of *σ* is set to 0.8.

#### 6 Inferring Higher-order Dependencies through *Matrix-based Rényi’s α Total Correlation*

Suppose now we have *n* ≥2 variables (*X*^1^, *X*^2^, …, *X*^*n*^) and a collection of *m* samples (Throughout this paper, we use superscript to denote variable index and subscript to denote sample index. For example, 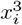 refers to the *i*-th sample from the 3rd variable.), i.e., 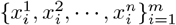, the *matrix-based Renyi’s α joint entropy* for *n* variables can be evaluated as [19]:

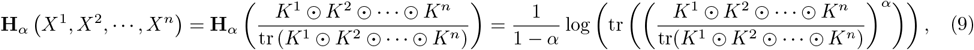

where 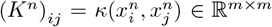 is a Gram matrix evaluated with *κ* for the *n*-th variable. The operator ⊙ is the Hadamard product.

The *Total Correlation* describes the dependence among *n* variables and can be considered as a non-negative generalization of the concept of *mutual information* from two parties to *n* parties. Let the definition of total correlation due Watanabe [13] be denoted as:

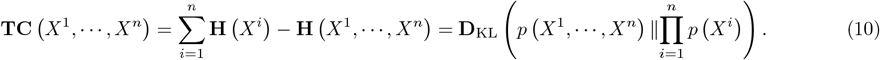

As seen above *Total Correlation* can also be equivalently expressed as the Kullback–Leibler divergence, **D**_KL_, between the joint probability density and the product of the marginal densities.

In order to estimate *Total Correlation* in a practical setting only from *m* samples 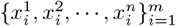, we convert Eqs. 10 to matrix-based *Renyi’s α entropy functional* based on Eqs. 4 and 9, which simplifies the estimation [19, 53] and enables a reformulation:

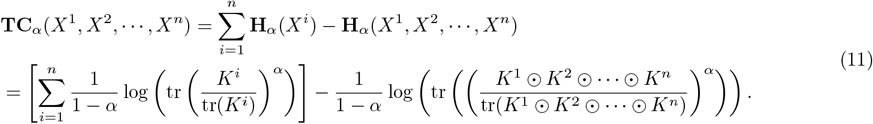

### E Estimating Pairwise and Beyond Pairwise Interactions in rsfMRI

#### Pairwise Interactions (Pearson Correlation & Mutual Information)

The rsfMRI signal comprising *n* ICNs obtained from scICA, denoted as *X*_*i*_ (1 ≤*i* ≤ *n* and *n* = 105), each *X*_*i*_ corresponds informally to a time instance *t*_*i*_ in a sequence {*t*_1_, *t*_2_, …, *t*_*T*_} with a constant time interval *t*. The pairwise interaction based on Pearson correlation can be estimated, where two ICNs that share substantial information are expected to exhibit a strong correlation, and vice versa. Meanwhile, the pairwise interaction based on mutual information can be estimated using Eq. 6. The mutual information measures the dependency between *X*_*i*_(*t*) and *X*_*j*_(*t*), and reaches its minimum value of zero when the two ICNs are independent.

#### 2. Beyond Pairwise Interactions (Total Correlation)

Estimating interactions beyond pairwise (***k*** *>* 2) among ***n*** = 105 ICNs, iterating over each set of indices used to obtain TC based on Eq. 11. If the beyond ICNs exhibit strong interactions, the TC will have a greater value, and vice versa. Additionally, TC is always positive.

### F Decomposing Triadic Interaction Tensors Using Canonical Polyadic Decomposition with Alternating Least Squares (CP-ALS)

Canonical Polyadic Decomposition (CP) is a popular method for analyzing and interpreting latent patterns in multidimensional data [30, 40]. One of the most widely used approaches for computing CP decomposition is the Alternating Least Squares (CP-ALS) method, which solves a series of linear least squares problems iteratively [30, 39, 40].

The goal of CP decomposition is to represent a tensor as the sum of rank-one components. For an *N*-mode ten-Sor 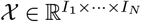, The n-mode matricization, or unfolding, of a 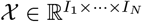, denoted 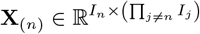, and the matrix **X**_(*n*)_ is formed so that the columns are the mode-*n* fibers of 𝒳. The rank-R CP decomposition of X is the approximation,

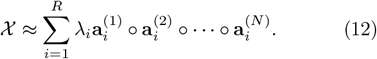

where tensors are denoted by boldface uppercase calligraphic letters, such as 𝒳, while vectors are denoted by boldface lowercase letters, such as **a**. The vectors 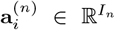 are unit vectors, with the weight vector *λ* ∈ℝ^*R*^, and ◦ denotes the outer product. The collection of all 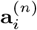 vectors for each mode is called a factor matrix, denoted as 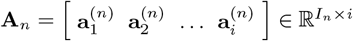.

To compute the CP decomposition, the CP-ALS algorithm solves a least squares problem for the matricized tensors **X**_(*n*)_ and 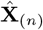 along each mode. For mode *n*, we fix every factor matrix except **Â**_(*n*)_ (here **Â**_*n*_ = **A**_*n*_ diag(***λ***)), and then solve for 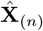. This process is repeated by alternating between modes until a termination criterion is met. We then solve the linear least squares problem,

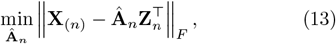

where 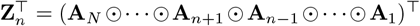 (denotes the Hadamard product), The linear least squares problem from Eq.13 is typically solved using the normal equations,

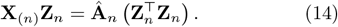

The coefficient matrix 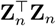 is computed efficiently. The desired factor matrix **A**_*n*_ is obtained by normalizing the columns of **Â**_*n*_ and updating the weight vector *λ*. The CP-ALS algorithm is already implemented in the MAT-LAB Tensor Toolbox (https://www.tensortoolbox.org/) [30].

In our study, we applied CP-ALS to triadic interaction tensors to uncover the hidden latent factors underlying complex interactions. This approach allows us to gain more precise insights into network interactions, revealing deeper patterns and structures that are often obscured within large volumes of triple interactions.

To evaluate the performance of the CP-ALS model fitting, we first convert the triadic interaction into a tensor object, enabling multi-dimensional array operations. The ground truth factors are represented as a CP decomposition with specified factor matrices and a weight vector, and the relative error between the full tensor and the data is computed. The true factors are then visualized using scatter plots for each mode (ICNs, ICNs, ICNs). Since these three models share the same pattern, we weight them together, and then present scatter plots of the latent factors for the 10 components separately. We then proceed to fit the CP decomposition using random initial guesses. A loop is executed for 30 iterations, and in each iteration, the CP decomposition of the data is estimated with a specified rank R=10. The relative error of each fit is computed, allowing us to monitor the progression of the model fitting and evaluate the accuracy of the decomposition.

To identify the top contributing ICNs in triple interactions, we selected the top three contributing ICNs in each component based on magnitude values. We also identified the top three contributing ICNs across all components. Later, we performed the same analysis for the control and SZ groups separately to identify any aberrant ICNs in the SZ group.

## V Data and Code Availability Statement

The data analyzed in this study cannot be shared without specific licenses. However, the dataset can be accessed upon request by contacting Prof.Vince D Calhoun.

The NeuroMark 2.2 templates are accessible on website (https://trendscenter.org/data/) and GitHub (https://github.com/trendscenter/gift/tree/master/GroupICAT/icatb/icatb_templates). The codes of the GICA, and MOO-ICAR have been integrated into the group ICA Toolbox (GIFT 4.0c, https://trendscenter.org/software/gift/). The MATLAB Tensor Toolbox is available at https://www.tensortoolbox.org/.

## Acknowledgments

This work was supported by NSF grant 2112455, and NIH grant R01MH123610. The authors thank anonymous reviewers for constructive feedback for improving the manuscript.

## References

[1] F. Battiston, G. Cencetti, I. Iacopini, V. Latora, M. Lucas, A. Patania, J.-G. Young, and G. Petri, Networks beyond pairwise interactions: Structure and dynamics, Physics Reports 874 (2020).

[2] J. D. Power, A. L. Cohen, S. M. Nelson, G. S. Wig, K. A. Barnes, J. A. Church, A. C. Vogel, T. O. Laumann, F. M. Miezin, B. L. Schlaggar, and S. E. Petersen, Functional network organization of the human brain, Neuron 72, 665 (2011).

[3] H.-J. Park and K. Friston, Structural and functional brain networks: from connections to cognition, Science 342, 1238411 (2013).

[4] B. T. Thomas Yeo, F. M. Krienen, J. Sepulcre, M. R. Sabuncu, D. Lashkari, M. Hollinshead, J. L. Roffman, J. W. Smoller, L. Zöllei, J. R. Polimeni, B. Fischl, H. Liu, and R. L. Buckner, The organization of the human cerebral cortex estimated by intrinsic functional connectivity, Journal of Neurophysiology 106, 1125 (2011).

[5] Q. Li, Functional connectivity inference from fmri data using multivariate information measures, Neural Networks 146, 85 (2022).

[6] M. Ashrafi and H. Soltanian-Zadeh, Multivariate gaussian copula mutual information to estimate functional connectivity with less random architecture, Entropy 24 (2022).

[7] Q. Li, G. Ver Steeg, and J. Malo, Functional connectivity via total correlation: Analytical results in visual areas, Neurocomputing 571, 127143 (2023).

[8] Q. Li, G. V. Steeg, S. Yu, and J. Malo, Functional connectome of the human brain with total correlation, Entropy 24, 1725 (2022).

[9] Q. Li, V. Calhoun, T. Pham, and A. Iraji, Exploring nonlinear dynamics in brain functionality through phase portraits and fuzzy recurrence plots, Chaos: An Interdisciplinary Journal of Nonlinear Science 34 (2024).

[10] R. Herzog, F. Rosas, R. Whelan, S. Fittipaldi, H. Santamaría-García, J. Cruzat, A. Birba, S. Moguilner, E. Tagliazucchi, P. Prado, and A. Ibanez, Genuine high-order interactions in brain networks and neurodegeneration, Neurobiology of Disease 175, 105918 (2022).

[11] M. Gatica, R. Cofré, P. A. Mediano, F. E. Rosas, P. Orio, I. Diez, S. P. Swinnen, and J. M. Cortes, High-order interdependencies in the aging brain, Brain connectivity 11, 734 (2021).

[12] Q. Xie, X. Zhang, I. Rekik, X. Chen, N. Mao, D. Shen, and F. Zhao, Constructing high-order functional connectivity network based on central moment features for diagnosis of autism spectrum disorder, PeerJ 9, e11692 (2021).

[13] S. Watanabe, Information theoretical analysis of multivariate correlation, IBM Journal of research and development 4, 66 (1960).

[14] T. S. Han, Nonnegative entropy measures of multivariate symmetric correlations, Information and Control 36, 133 (1978).

[15] L. Xiao, J. Wang, P. H. Kassani, Y. Zhang, Y. Bai, J. M. Stephen, T. W. Wilson, V. D. Calhoun, and Y. ping Wang, Multi-hypergraph learning-based brain functional connectivity analysis in fmri data, IEEE Transactions on Medical Imaging 39, 1746 (2019).

[16] F. A. N. Santos, E. P. Raposo, M. D. Coutinho-Filho, M. Copelli, C. J. Stam, and L. Douw, Topological phase transitions in functional brain networks, Phys. Rev. E 100, 032414 (2019).

[17] E. Zampieri, G. Moreni, C. Vriend, L. Douw, and F. Santos, A hands-on tutorial on network and topological neuroscience, Brain Structure and Function 227 (2022).

[18] Q. Li, S. Yu, K. H. Madsen, V. D. Calhoun, and A. Iraji, Higher-order organization in the human brain from matrix-based rényi’s entropy, in 2023 IEEE International Conference on Acoustics, Speech, and Signal Processing Workshops (ICASSPW) (2023) pp. 1–5.

[19] S. Yu, L. G. S. Giraldo, R. Jenssen, and J. C. Principe, Multivariate extension of matrix-based renyi’s α-order entropy functional, IEEE transactions on pattern analysis and machine intelligence 42, 2960 (2019).

[20] V. Laparra, G. Camps-Valls, and J. Malo, Iterative Gaussianization: From ICA to random rotations, IEEE Transactions on Neural Networks 22, 537 (2011).

[21] V. Laparra, J. E. Johnson, G. Camps-Valls, R. Santos-Rodríguez, and J. Malo, Estimating information theoretic measures via multidimensional gaussianization, IEEE Transactions on Pattern Analysis and Machine Intelligence 47, 1293 (2025).

[22] Q. Li, V. Calhoun, A. R. Ballem, S. Yu, J. Malo, and A. Iraji, Aberrant high-order dependencies in schizophrenia resting-state functional MRI networks, in NeurIPS 2023 workshop: Information-Theoretic Principles in Cognitive Systems (2023).

[23] V. D. Calhoun, T. Adalı, G. D. Pearlson, and J. J. Pekar, A method for making group inferences from functional mri data using independent component analysis, Human Brain Mapping 14 (2001).

[24] E. Allen, E. Erhardt, E. Damaraju, W. Gruner, J. Segall, R. Silva, M. Havlicek, S. Rachakonda, J. Fries, R. Kalyanam, A. Michael, A. Caprihan, J. Turner, T. Eichele, S. Adelsheim, A. Bryan, J. Bustillo, V. Clark, S. Ewing, and V. Calhoun, A baseline for the multivariate comparison of resting-state networks, Frontiers in systems neuroscience 5, 2 (2011).

[25] Y. Du, Z. Fu, J. Sui, S. Gao, Y. Xing, D. Lin, M. Salman, A. Abrol, M. A. Rahaman, J. Chen, P. Kochunov, E. Osuch, and V. Calhoun, Neuromark: An automated and adaptive ica based pipeline to identify reproducible fmri markers of brain disorders, NeuroImage: Clinical 28, 102375 (2020).

[26] H. Morioka, V. Calhoun, and A. Hyvärinen, Nonlinear ica of fmri reveals primitive temporal structures linked to rest, task, and behavioral traits, NeuroImage 218, 116989 (2020).

[27] X. Liu and J. H. Duyn, Time-varying functional network information extracted from brief instances of spontaneous brain activity, Proceedings of the National Academy of Sciences 110, 4392 (2013).

[28] A. Iraji, Z. Fu, A. Faghiri, M. Duda, J. Chen, S. Rachakonda, T. DeRamus, P. Kochunov, B. Adhikari, A. Belger, J. Ford, D. Mathalon, G. Pearlson, S. Potkin, A. Preda, J. Turner, T. Erp, J. Bustillo, K. Yang, and V. Calhoun, Identifying canonical and replicable multiscale intrinsic connectivity networks in 100k+ restingstate fmri datasets, Human Brain Mapping 44 (2023).

[29] K. Jensen, J. Turner, V. Calhoun, and A. Iraji, Addressing inconsistency in functional neuroimaging: A replicable data-driven multi-scale functional atlas for canonical brain networks, biorxiv (2024).

[30] B. W. Bader, T. G. Kolda, et al., Tensor toolbox for matlab, version 3.6, https://www.tensortoolbox.org (2023).

[31] V. Menon, 20 years of the default mode network: A review and synthesis, Neuron 111 (2023).

[32] J. S. Damoiseaux, S. A. R. B. Rombouts, F. Barkhof, P. Scheltens, C. J. Stam, S. M. Smith, and C. F. Beckmann, Consistent resting-state networks across healthy subjects, Proceedings of the National Academy of Sciences 103, 13848 (2006).

[33] V. Calhoun, J. Sui, K. Kiehl, J. Turner, E. Allen, and G. Pearlson, Exploring the psychosis functional connectome: Aberrant intrinsic networks in schizophrenia and bipolar disorder, Frontiers in psychiatry / Frontiers Research Foundation 2, 75 (2011).

[34] E. Dellen, C. Börner-Schröder, M. Schutte, S. Montfort, L. Abramovic, M. Boks, W. Cahn, N. Haren, R. Mandl, C. Stam, and I. Sommer, Functional brain networks in the schizophrenia spectrum and bipolar disorder with psychosis, npj Schizophrenia 6, 22 (2020).

[35] M.-E. Lynall, D. Bassett, R. Kerwin, P. McKenna, M. Kitzbichler, U. Müller-Sedgwick, and E. Bullmore, Functional connectivity and brain networks in schizophrenia, The Journal of neuroscience : the official journal of the Society for Neuroscience 30, 9477 (2010).

[36] L. Torres, A. S. Blevins, D. Bassett, and T. Eliassi-Rad, The why, how, and when of representations for complex systems, SIAM Review 63, 435 (2021).

[37] E. Allen, E. Damaraju, S. Plis, E. Erhardt, T. Eichele, and V. Calhoun, Tracking whole-brain connectivity dynamics in the resting state, Cerebral cortex (New York, N.Y. : 1991) (2012).

[38] D. Vidaurre, S. M. Smith, and M. W. Woolrich, Brain network dynamics are hierarchically organized in time, Proceedings of the National Academy of Sciences of the United States of America 114, 12827 (2017).

[39] B. W. Bader and T. G. Kolda, Matlab tensor classes for fast algorithm prototyping, ACM Transactions on Mathematical Software - TOMS (2004).

[40] B. W. Bader and T. G. Kolda, Efficient matlab computations with sparse and factored tensors, SIAM J. Scientific Computing 30, 205 (2007).

[41] C. A. Tamminga, E. I. Ivleva, M. S. Keshavan, G. D. Pearlson, B. A. Clementz, B. Witte, D. W. Morris, J. Bishop, G. K. Thaker, and J. A. Sweeney, Clinical phenotypes of psychosis in the bipolar-schizophrenia network on intermediate phenotypes (b-snip), American Journal of psychiatry 170, 1263 (2013).

[42] S. Meda, G. Ruaño, A. Windemuth, K. O’Neil, C. Berwise, S. Dunn, L. Boccaccio, B. Narayanan, M. Kocherla, E. Sprooten, M. Keshavan, C. Tamminga, J. Sweeney, B. Clementz, V. Calhoun, and G. Pearlson, Multivariate analysis reveals genetic associations of the resting default mode network in psychotic bipolar disorder and schizophrenia, Proceedings of the National Academy of Sciences of the United States of America 111 (2014).

[43] A. Iraji, A. Faghiri, N. Lewis, Z. Fu, S. Rachakonda, and V. Calhoun, Tools of the trade: Estimating time-varying connectivity patterns from fmri data, Social cognitive and affective neuroscience 16 (2020).

[44] X. Meng, A. Iraji, Z. Fu, P. Kochunov, A. Belger, J. M. Ford, S. McEwen, D. H. Mathalon, B. A. Mueller, G. Pearlson, S. G. Potkin, A. Preda, J. Turner, T. G. van Erp, J. Sui, and V. D. Calhoun, Multi-model order spatially constrained ica reveals highly replicable group differences and consistent predictive results from resting data: A large n fmri schizophrenia study, NeuroImage: Clinical 38, 103434 (2023).

[45] E. Erhardt, S. Rachakonda, E. Bedrick, E. Allen, T. Adali, and V. Calhoun, Comparison of multi-subject ica methods for analysis of fmri data, Human brain mapping 32, 2075 (2011).

[46] Q.-H. Lin, J. Liu, Y.-R. Zheng, H. Liang, and V. Calhoun, Semiblind spatial ica of fmri using spatial constraints, Human brain mapping 31, 1076 (2009).

[47] W. Lu and J. Rajapakse, Approach and applications of constrained ica, IEEE Transactions on Neural Networks 16, 203 (2005).

[48] C. E. Shannon, A mathematical theory of communication, The Bell system technical journal 27, 379 (1948).

[49] A. Rényi, On measures of entropy and information, in Proceedings of the Fourth Berkeley Symposium on Mathematical Statistics and Probability, Volume 1: Contributions to the Theory of Statistics, Vol. 4 (University of California Press, 1961) pp. 547–562.

[50] T. M. Cover and J. A. Thomas, Information theory and statistics, Elements of information theory 1, 279 (1991).

[51] L. G. S. Giraldo, M. Rao, and J. C. Principe, Measures of entropy from data using infinitely divisible kernels, IEEE Transactions on Information Theory 61, 535 (2014).

[52] D. J. Henderson and C. F. Parmeter, Normal reference bandwidths for the general order, multivariate kernel density derivative estimator, Statistics & Probability Letters 82, 2198 (2012).

[53] S. Yu, F. Alesiani, X. Yu, R. Jenssen, and J. Principe, Measuring dependence with matrix-based entropy functional, in Proceedings of the AAAI Conference on Artificial Intelligence, Vol. 35 (2021) pp. 10781–10789.

